# dbOTU3: A new implementation of distribution-based OTU calling

**DOI:** 10.1101/076927

**Authors:** Scott Olesen, Claire Duvallet, Eric Alm

## Abstract

Distribution-based operational taxonomic unit-calling (dbOTU) improves on other approaches by incorporating information about the input sequences’ distribution across samples. Previous implementations of dbOTU presented challenges for users. Here we introduce and evaluate a new implementation of dbOTU that is faster and more user-friendly. We show that this new implementation has theoretical and practical improvements over previous implementations of dbOTU, making the algorithm more accessible to microbial ecology and biomedical researchers.

## 2 Introduction

Preheim *et al*. [1] formulated the distribution-based OTU-calling (dbOTU) algorithm, an extremely accurate algorithm for grouping DNA sequences from microbial communities into operational taxonomic units (OTUs) for ecological or biomedical research. Unlike most OTU-calling approaches, which group sequences based only on the similarities of the sequences themselves, this algorithm also uses information about the distribution of sequences across samples. This allows dbOTU to distinguish ecologically-distinct but sequence-similar organisms or populations.

### 2.1 The algorithm

#### 2.1.1 Motivation

The algorithm aims to separate genetically-similar sequences that appear to be ecologically distinct (or, conversely, to join less-genetically-similar sequences that appear to be ecologically identical). For example, if two sequences differ by only one nucleotide, an OTU-calling algorithm would likely group those two sequences into the same OTU. However, if the two sequences never appeared together in the same sample, an observer would probably conclude that that one nucleotide difference corresponds to two distinct groups of organisms, one which lives in one group of samples, the other living in the other.

Conversely, if two sequences differed by a few nucleotides, an OTU-calling algorithm would probably place two sequences into different OTUs. However, if the two sequences appeared in the same ratio in all samples (e.g., sequence 2 was always almost exactly ten times less abundant than sequence 1), an observer might conclude that the second sequence was either sequencing error or a member of the same ecological population as the first sequence.

#### 2.1.2 Mechanics

The original workflow was:

1. Process 16S data up to dereplicated sequences.
2. Create a table of sequence counts showing the number of times each sequence appears in each sample.
3. Align the dereplicated sequences. Using the alignment, make a phylogenetic tree and a “distance matrix” showing the genetic dissimilarity between sequences.
4. Feed the matrix and the table of sequence counts into the algorithm proper, which groups the sequences into OTUs.

In outline, step 4 meant:

1. Make the most abundant sequence an OTU.
2. For each sequence (in order of decreasing abundance), find the set of OTUs that meet “abundance” and “genetic” criteria. The abundance criterion requires that the candidate sequence be some fold less abundant than the OTU (e.g., so that it can be considered sequencing error). The genetic criterion requires that the candidate sequence be sufficiently similar to the OTU’s sequence (e.g., so that it can be considered sequencing error or part of the same population of organisms).
3. If no OTUs meet these two criteria, make the candidate sequence into a new OTU.
4. If OTUs do meet these criteria, then, starting with the most closely-genetically-related OTU, check if the candidate sequence is distributed differently among the samples than that OTU. If the distributions are sufficiently similar, merge the candidate sequence into that OTU. Specifically, add the candidate sequence’s counts across samples to the OTU’s counts.
5. If the candidate sequence does not have a distribution across sample sufficiently similar to an existing OTU, then make this sequence a new OTU.
6. Move on to the next candidate sequence.

Note that an OTU’s counts change every time a candidate sequence is merged into that OTU, but an OTU’s sequence never changes. In other words, an OTU’s candidate sequence is the sequence of its most abundant member.

### 2.2 Previous implementations

The dbOTU algorithm has been implemented twice. Here we will introduce a third implementation. The implementations vary in terms of:

- the exact input files they require,
- how they evaluate the genetic (i.e., sequence similarity) criterion,
- how they evaluate the distribution (i.e., ecological similarity) criterion, and
- the details of the software itself.

These differences are summarized in Table 1.

**Table 1:** Comparison of the dbOTU implementations. *In the first two implementations, the dissimilarity between two sequences was the minimum of the dissimilarities of the aligned and unaligned sequences.

| Imple-<br>mentation | Programming<br>languages | Required input | Genetic test | Distribution<br>test |
| --- | --- | --- | --- | --- |
| 1 | Perl, R, shell | Aligned sequences,<br>matrix of genetic distances,<br>sequence count table | Input<br>dissimilarities* | Simulated<br>$\chi^2$ test |
| 2 | Python 2,<br>Perl, R | Aligned sequences,<br>sequence count table | Proportion of<br>mismatched sites* | Simulated<br>$\chi^2$ test |
| 3 | Python 3 | Unaligned sequences,<br>sequence count table | Levenshtein<br>edit distance | Likelihood-ratio test |

#### 2.2.1 The first implementation

The original implementation^1^, coded in Perl and shell scripts, took a matrix of genetic dissimilarities as input and used a *χ*^2^ of independence as the distribution criterion.

In the original publication, the Jukes-Cantor distance was used as the genetic dissimilarity metric. The Jukes-Cantor distance is 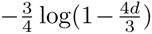, where *d* is the proportion of positions that differ (i.e., the number of mismatches in the aligned sequences divided by the length of the aligned sequences, ignoring gaps).

This implementation also required aligned sequences as input. In the original publication, sequences were aligned using the `align.seqs` command in `mothur` [2], which implemented the NAST alignment algorithm [3], and the Jukes-Cantor distances were computed using the `-makematrix` option in Fast-Tree [4]. The resulting matrix of distances was used as input to the software. The genetic criterion was actually articulated as the minimum of the aligned and unaligned Jukes-Cantor distances, which was a work-around for the fact that using the aligned sequences sometimes led to a greater distance between two sequences than would be computed by just comparing the unaligned sequences.

In this implementation, the distribution criterion was evaluated using the *χ*^2^ test of independence as implemented in the `chisq.test` function in `R` [5], called in a separate process from a Perl script. Many of the comparisons involved sequences with small numbers of counts, for which the asymptotic (i.e., commonly-used) calculation of the *p*-value of a *χ*^2^ test is not accurate. This implementation therefore used a simulated *p*-value, available through the `R` command’s `simulate.p.value` option. This empirical calculation required many simulated contingency tables, which was expensive.

#### 2.2.2 The second implementation

The second implementation^2^, coded in Python 2 and interfaced with R using `r2py`^3^, also requires aligned sequences as input uses the simple metric *d* (as defined above), rather than the Jukes-Cantor distance. Rather than taking a matrix of pairwise distances, the dissimilarities between sequences were computed only as necessary, which eliminated the requirement to store *N*^2^ numbers in memory, where *N* is the number of sequences to be clustered. Like the first implementation, this one used the minimum of the aligned and unaligned sequences.

Like the first implementation, this one used R’s `chisq.test`, but this time called via `r2py` from the Python script. This removed the need for temporary files, but it was still slow and required both R and Python.

### 2.3 The genetic criterion

A critical piece of the dbOTU algorithm is determining which sequences are sufficiently genetically dissimilar that they belong in different OTUs regardless of their distribution across samples.

The first two implementations used sequence dissimilarities that relied on alignments. The alignments were apparently not foolproof, as evidenced by the need for using the minimum of the distances between the aligned and unaligned pairs of sequences.

### 2.4 The distribution criterion

The first two implementations tested for the independence of the candidate sequence’s and OTU’s distribution across samples using an empirical *p*-value. When the number of counts in the candidate sequence and OTU being compared were large, the asymptotic *p*-value for the *χ*^2^ test was valid, which made it easy to compute. However, the *χ*^2^ test is fairly sensitive: it often concludes that a candidate sequence and OTU are distributed differently when inspection suggests that the differences between the two distributions are due to noise and not true ecological differences.

For example, if a candidate sequence and an OTU had relative abundances of 50% and 55% respectively, each supported by ten thousand counts, then the *χ*^2^ will register the two as different, even though a human observer could easily conclude that the 5% different in relative abundance was due to noise inherent in the sampling, library preparation, or sequencing procedures. Thus, using the *χ*^2^ test will sometimes prevent sequences that should be merged from being merged.

#### 2.4.1 This implementation

This implementation, dbOTU3, aims to improve speed and ease of use. It is written in pure Python 3. Rather than aligning sequences, this implementation uses the Levenshtein edit distance (from the python-Levenshtein package^4^) as an approximation for the sequence dissimilarity. Rather than using an empirical *χ*^2^ test, this implementation uses a likelihood ratio test. The merit of these choices are discussed in the Results.

## 3 Methods

### 3.1 New genetic and distribution criteria

This implementation evaluates the genetic criterion using the Levenshtein edit distance, i.e., a candidate sequence will not be merged into an OTU if 2*E/*(*ℓ*_1_ + *ℓ*_2_) is greater than some threshold, where *E* is the Levenshtein edit distance, *ℓ*_1_ is the length of the candidate sequence, and *ℓ*_2_ is the length of the OTU’s sequence. As shown in the Results, this metric is a good approximation of the proportion *d* of mismatched sites in an alignment.

This implementation evaluates the distribution criterion using a likelihood-ratio test. Define 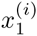 be the number of counts that the OTU has in sample *i* and 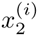 be the number of counts the candidate sequence has. Define also 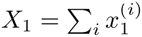 and similarly *X*_2_.

The alternative hypothesis for this test is that the OTU and candidate sequence are distributed “differently”, that is, that each of the 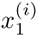 and 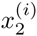 are draft from different random variables. Specifically, the alternative hypothesis is

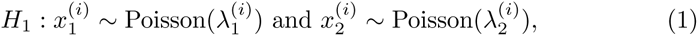

where there are no constraints on the Poisson parameters.

The null model asserts that the OTU and candidate sequence are distributed “the same”, that is, that the candidate sequence’s counts in each sample is drawn from a Poisson random variable whose parameter is proportional to the parameter of the OTU’s Poisson variables, where the constant of proportionality is the same across samples. Specifically, the null model is

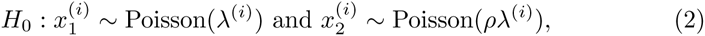

We expect that, because the candidate sequence is overall less abundant than the OTU, 0 *< ρ <* 1.

Asserting maximum likelihood for each model shows that

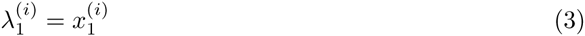

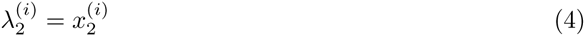

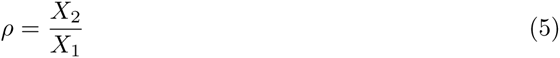

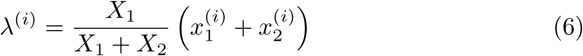

so that the test statistic 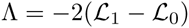 is

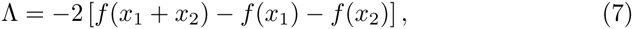

where 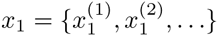 and

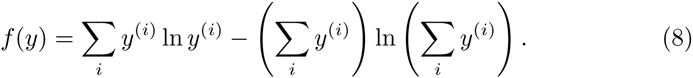

### 3.2 Accuracy of new implementation

To evaluate the performance of the new implementation, we compared the results of calling OTUs with this dbOTU implementation, the second (previous) dbOTU implementation, and with UPARSE [6]. We used the Turnbaugh mock community data set analyzed in the original publication [7].

To prepare the data for input into the OTU callers, we first downloaded the data^5^, which included:

- `Mock clean.fna:` quality-screened sequences from all 6 mock communities
- `Mock nochimeras.fna:` quality-screened, de-noised, and non-chimeric sequences.

We trimmed all sequences in `Mock nochimeras.fna` to 187 nucleotides, the length of the shortest sequence in that file. To align the sequences in `Mock clean.fna` with those in `Mock nochimeras.fna`, we trimmed the first 14 nucleotides from each sequence in `Mock clean.fna` and then trimmed the remaining sequences to 187 nucleotides.

To generate a table of sequence counts, we dereplicated the unique trimmed sequences from `Mock nochimeras.fna`. For each sequence in `Mock clean.fna`, we checked if that sequence appeared among the unique sequences. If so, we counted it as present in the sample corresponding to the metadata for that sequence.

To generate the aligned sequences required for the previous dbOTU implementation, we aligned the unique sequences using QIIME’s `align seqs.py` with default parameters [8].

Both dbOTU implementations were run using a genetic dissimilarity threshold of 0.1 and using *p* = 0.001 as the distribution test threshold. UPARSE OTUs were called using `usearch -cluster otus` and the same list of unique sequences (but with the summary “size” information required by `usearch`) at similarity thresholds of 95%, 97%, and 100%. Notably, UPARSE includes chimera checking, which identified chimeras among the sequences from `Mock nochimeras.fna.` The specific sequences that UPARSE identified as chimeras depended on the clustering similarity threshold.

We compared the results of the OTU callers with the true composition of each mock community sample (Table S3 in Turnbaugh et al. [7]). To link the sequence data with the true mock community composition, which is expressed in terms of the abundances of input isolates, we identified, for each dereplicated sequence, the most genetically-similar reference sequence in `MockIsolatesV2.fna, specifically`, the one with the smallest dissimilarity *d* to the query sequence. To compare compositions, we combined the abundance data from all the dereplicated sequences that mapped to the same species.

### 3.3 Speed of the genetic tests

To benchmark the speed of the new genetic test, dissimilarity between the first two sequences in the list of dereplicated sequences from the mock community was computed 10 times using each of three alignment methods: the new Levenshtein-based method, Biopython’s `pairwise2.align.globalxx` [9], and an external call to Clustal Omega [10].

### 3.4 Accuracy of the genetic tests

To evaluate the performance of the new Levenshtein-based genetic dissimilarity metric, we compared the dissimilarities computed by the new metric, the previously-implemented unaligned and aligned dissimilarity metrics, and a gold standard: a pairwise alignment of the two sequences using Clustal Omega. We computed genetic dissimilarities for all pairs of sequences for which the genetic test was evaluated during OTU calling on the mock data set.

### 3.5 Accuracy of the distribution tests

To evaluate the performance of the new likelihood-ratio test, we compared the *p*-values computed by the new test against the simulated *χ*^2^ test used in earlier implementations. 10^7^ simulations were used to compute the *p*-value for that test. For comparison, the same computations were performed using the asymptotic *χ*^2^ test. Accuracies were quantified *F*-scores (considering the simulated *χ*^2^ test as the truth).

## 4 Results

### 4.1 The new implementation performs similarly to the previous one

This dbOTU implementation and the previous implementation yield nearly identical species-level compositions (Figure 1). Both dbOTU implementations performed similarly to 100% clustering with UPARSE but less similarly to other clustering thresholds with UPARSE.

**Figure 1:**
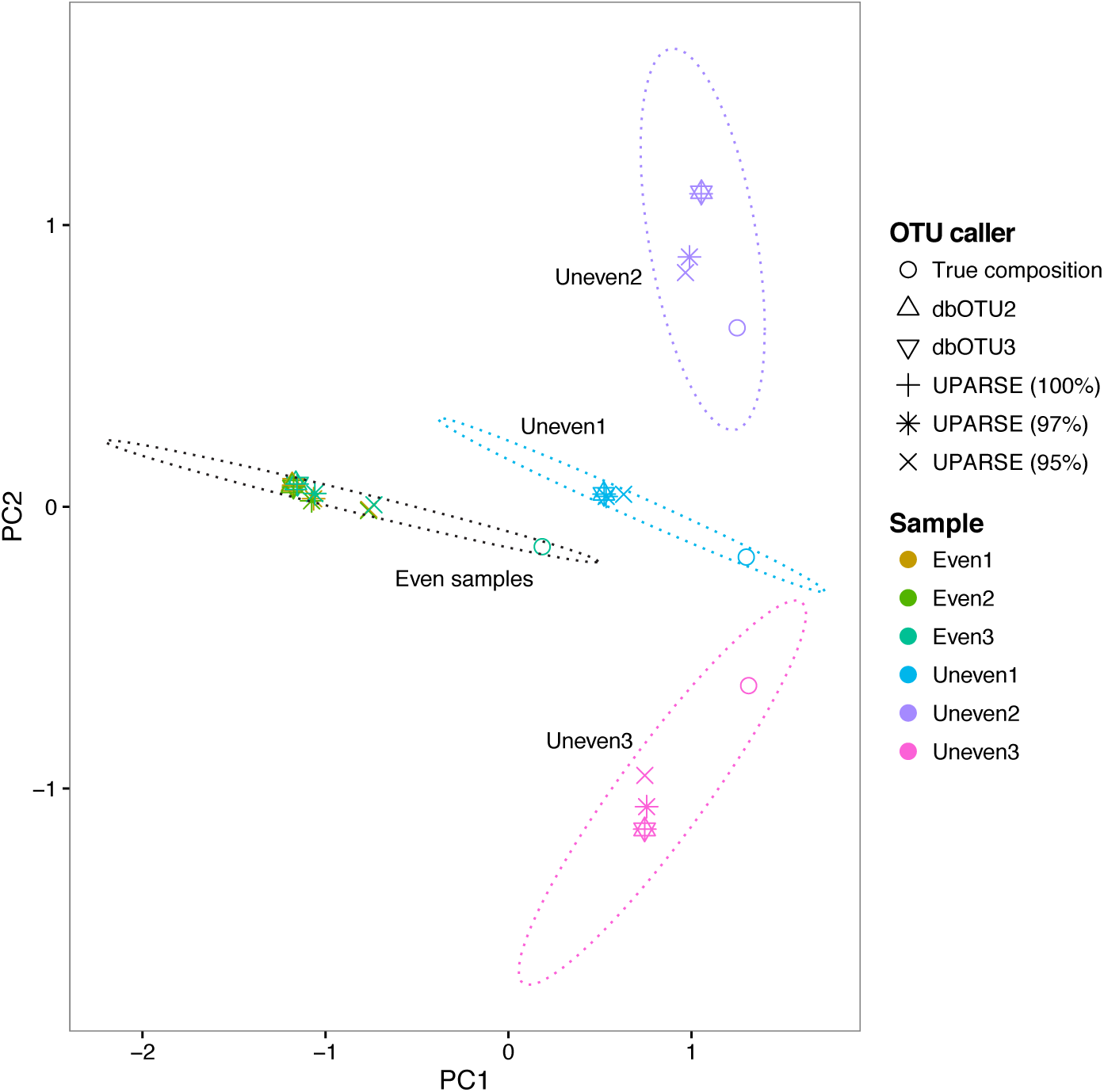
This implementation (dbOTU3) produces nearly identical results with the previous implementation (dbOTU2) as visualized in a principal coordinate analysis ordination plot, communities resulting from analysis of the mock community data with various OTU callers. (The two triangles representing dbOTU2 and dbOTU3 always appear on top of one another, making a six-pointed triangle.) The “true composition” is the community composition expected based on how the communities were constructed. The principal components were computed using a matrix of the square roots of the Jensen-Shannon divergence between each pair of computed community compositions.

### 4.2 The Levenshtein-based dissimilarity is fast and performs as well as the previous metric

The Levenshtein-based genetic dissimilarity was much faster than either in-Python alignments made using Biopython’s alignment or out-of-Python alignments using Clustal Omega (Table 2).

**Table 2:** Benchmarks for the speed of the dissimilarity metric computations relative to the Levenshtein-based metric.

| Metric | Relative time |
| --- | --- |
| Levenshtein-based | 1.0 |
| Clustal Omega | 210.4 |
| Biopython | 3127.2 |

The correlations between the gold standard (the pairwise-alignments made with Clustal Omega) and the other genetic dissimilarity metrics are shown in Table 3. The new Levenshtein-based metric performs comparably (or even slightly better than) the minimum of the aligned and unaligned *d* dissimilarities, which was used for the previous implementations.

**Table 3:** Correlation coefficients between genetic dissimilarities computed by a pairwise alignment using Clustal Omega and other metrics. *d* is the proportion of mismatched sites.

| Metric | Correlation (%) | 95% confidence interval (%) |
| --- | --- | --- |
| Levenshtein-based | 82.8 | 72.5–89.5 |
| Minimum $d$ | 80.5 | 69.1–88.0 |
| Aligned $d$ | 49.5 | 27.1–66.8 |
| Unaligned $d$ | 40.8 | 16.8–60.3 |

### 4.3 The likelihood-ratio test mostly reproduces the previous distribution criterion

When calling OTUs on the mock data set, the new implementation’s distribution criterion performed similarly to the simulated *χ*^2^ test. Of 58 comparisons, the simulated *χ*^2^ determined that the candidate sequence and OTU in question were differently distributed in 25 cases. The likelihood-ratio test concurred in those 25 cases. In 4 cases, the simulated *χ*^2^ test identified the candidate sequence and OTU as similarly distributed but the likelihood-ratio test identified them as differently distributed. Those cases are shown in Table 4. This performance (25 true positives and 4 false positives) corresponds to an accuracy of *F*_1_ = 93%, while the asymptotic *χ*^2^ achieved an accuracy of only *F*_1_ = 79% (25 true positives and 13 false positives).

**Table 4:** The four cases for which the simulated *χ*^2^ test determines that the OTU (top row) and candidate sequence (bottom row) are identically distributed but the likelihood-ratio test determines that they are differently distributed (both with respect to the threshold *p* = 0.001).

| IDs | Counts |  |  |  |  |  |
| --- | --- | --- | --- | --- | --- | --- |
|  | Even1 | Even2 | Even3 | Uneven1 | Uneven2 | Uneven3 |
| seq84 | 462 | 398 | 622 | 17 | 96 | 1 |
| seq40 | 33 | 25 | 26 | 4 | 0 | 1 |
| seq15 | 138 | 129 | 163 | 92 | 257 | 14 |
| seq45 | 15 | 11 | 28 | 1 | 13 | 1 |
| seq31 | 193 | 151 | 254 | 11 | 497 | 2 |
| seq45 | 15 | 11 | 28 | 1 | 13 | 1 |
| seq25 | 172 | 145 | 279 | 7 | 512 | 5 |
| seq45 | 15 | 11 | 28 | 1 | 13 | 1 |

Up to this point, the *p*-value threshold for the likelihood-ratio test was fixed at the same value as was used for the *χ*^2^ test. To check if the likelihood-ratio test would perform better when using a different threshold, we varied the likelihood-ratio test’s threshold, keeping the *χ*^2^ test’s threshold fixed, and computed the new test’s accuracy (Table 5). The likelihood-ratio test performs best, relative to the *χ*^2^ test, when its *p*-value threshold is about ten-fold smaller.

## 5 Discussion

This implementation is overall more user-friendly than previous implementations. The software itself is faster, and the base codebase is smaller. The main codebase is all in a single file (275 lines) which is supported by online documentation and unit tests. The dependencies for running the code are simpler: it requires only a few Python packages. The input files for this implementation are easier for the user to prepare because they do not require an alignment step.

**Table 5:** The likelihood-ratio test perfectly reproduces the results of the *χ*^2^ test for a smaller *p*-value threshold (~0.0001). In these comparisons, the *χ*^2^ test’s *p*-value threshold was fixed at 0.001, and the likelihood-ratio test’s threshold was adjusted to intermediate values appearing in the list of *p*-values the test computed. The *χ*^2^ test delivered 25 positives.

| Threshold<br>$p$-value | True<br>positives | All<br>positives | $F_1$<br>(%) |
| --- | --- | --- | --- |
| 0.000000100 | 21 | 21 | 91.3 |
| 0.000000385 | 21 | 21 | 91.3 |
| 0.000000396 | 22 | 22 | 93.6 |
| 0.000001955 | 23 | 23 | 95.8 |
| 0.000004453 | 24 | 24 | 98.0 |
| 0.000244656 | 25 | 25 | 100.0 |
| 0.000351707 | 25 | 26 | 98.0 |
| 0.000352663 | 25 | 27 | 96.2 |
| 0.000516666 | 25 | 28 | 94.3 |
| 0.001740844 | 25 | 29 | 92.6 |
| 0.002082822 | 25 | 30 | 90.9 |
| 0.002162794 | 25 | 31 | 89.3 |
| 0.007798747 | 25 | 32 | 87.7 |
| 0.009193968 | 25 | 33 | 86.2 |
| 0.013793169 | 25 | 34 | 84.7 |
| 0.014450072 | 25 | 35 | 83.3 |
| 0.015746250 | 25 | 36 | 82.0 |
| 0.017184692 | 25 | 37 | 80.6 |
| 0.033304330 | 25 | 38 | 79.4 |
| 0.069550045 | 25 | 39 | 78.1 |
| 0.074798381 | 25 | 40 | 76.9 |
| 0.094045427 | 25 | 41 | 75.8 |

Aside from software, this implementation has two main differences from the previous implementations. The first is the genetic dissimilarity criterion. Ideally, the genetic dissimilarity metric would be computed by a pairwise alignment of sequences. Unfortunately, there is no efficient implementation of this kind of alignment in pure Python. We found that calling Clustal Omega as a separate process was more than a hundred times slower than computed the Levenshtein distance. If this kind of computation is efficiently implemented in Python (perhaps in `scikit-bio`^6^), the Levenshtein-based metric should be replaced. In the meantime, the Levenshtein-based dissimilarity is efficient and well-supported.

The second major difference from previous implementations is the distribution criterion. In previous implementations, the simulated *χ*^2^ test, which is computationally demanding, was used in place of the asymptotic *χ*^2^ test, which is not computationally demanding, because the asymptotic *χ*^2^ test is not accurate when some cells have low counts (i.e., the candidate sequence is overall rare or is absent or nearly absent from some samples). This implementation used a likelihood-ratio test that performed similarly to the simulated *χ*^2^ test when using the same *p*-value threshold. When using the same threshold, the few cases in which the likelihood-ratio test and the simulated *χ*^2^ test divergence do not share any obvious pattern (Table 4). Adjusting the threshold for the likelihood-ratio test to ten-fold smaller perfectly reproduced the criterion emerging from the *χ*^2^ test. We therefore recommend that users migrating from dbOTU1 or 2 to dbOTU3 adjust their *p*-value threshold similarly.

In the analysis of the mock community data, these differences in implementation of the genetic and distribution criteria had a negligible effect on the resulting inferred community compositions. Therefore, the degree to which this implementation’s results did not recapitulate the “true” compositions is addressed by the original publication, which rigorously showed that the dbOTU algorithm was more accurate than alternative methods.

## 6 Acknowledgements

We thank Sarah Preheim for reviewing and suggesting improvements to this manuscript.

## Footnotes

1 https://github.com/spacocha/Distribution-based-clustering

2 https://github.com/spacocha/dbOTUcaller

3 http://rpy2.bitbucket.org

4 https://github.com/ztane/python-Levenshtein

5 https://gordonlab.wustl.edu/TurnbaughSE_2_10/PNAS_2010.html

6 http://scikit-bio.org

## References

[1] S. P. Preheim, A. R. Perrotta, A. M. Martin-Platero, A. Gupta, and E. J. Alm. Distribution-based clustering: Using ecology to refine the operational taxonomic unit. Appl Environ Microbiol, 79(21):6593–6603, 2013. doi: 10.1128/aem.00342-13. URL http://dx.doi.org/10.1128/AEM.00342-13.

[2] P. D. Schloss, S. L. Westcott, T. Ryabin, J. R. Hall, M. Hartmann, E. B. Hollister, R. A. Lesniewski, B. B. Oakley, D. H. Parks, C. J. Robinson, J. W. Sahl, B. Stres, G. G. Thallinger, D. J. Van Horn, and C. F. Weber. Introducing mothur: Open-source, platform-independent, community-supported software for describing and comparing microbial communities. Appl Environ Microbiol, 75(23):7537–7541, 2009. doi: 10.1128/aem.01541-09. URL http://dx.doi.org/10.1128/AEM.01541-09.

[3] T. Z. DeSantis, P. Hugenholtz, K. Keller, E. L. Brodie, N. Larsen, Y. M. Piceno, R. Phan, and G. L. Andersen. NAST: a multiple sequence alignment server for comparative analysis of 16s rRNA genes. Nucleic Acids Res, 34(Web Server):W394–W399, 2006. doi: 10.1093/nar/gkl244. URL http://dx.doi.org/10.1093/nar/gkl244.

[4] Morgan N. Price, Paramvir S. Dehal, and Adam P. Arkin. FastTree 2 – approximately maximum-likelihood trees for large alignments. PLoS One, 5(3):e9490, 2010. doi: 10.1371/journal.pone.0009490. URL http://dx.doi.org/10.1371/journal.pone.0009490.

[5] R Core Team. R: A Language and Environment for Statistical Computing. R Foundation for Statistical Computing, Vienna, Austria, 2016. URL https://www.R-project.org/.

[6] Robert C Edgar. UPARSE: highly accurate OTU sequences from microbial amplicon reads. Nat Methods, 10(10):996–998, 2013. doi: 10.1038/nmeth.2604. URL http://dx.doi.org/10.1038/nmeth.2604.

[7] P. J. Turnbaugh, C. Quince, J. J. Faith, A. C. McHardy, T. Yatsunenko, F. Niazi, J. Affourtit, M. Egholm, B. Henrissat, R. Knight, and J. Gordon. Organismal, genetic, and transcriptional variation in the deeply sequenced gut microbiomes of identical twins. Proc Natl Acad Sci USA, 107(16):7503–7508, 2010. doi: 10.1073/pnas.1002355107. URL http://dx.doi.org/10.1073/pnas.1002355107.

[8] J.G. Caporaso, J. Kuczynski, J. Stombaugh, K. Bittinger, F.D. Bushman, E.K. Costello, N. Fierer, A.G. Pena, J.K. Goodrich, J.I. Gordon, and G.A. Huttley. QIIME allows analysis of high-throughput community sequencing data. Nature, 7(5):335–336, 2010. doi: doi:10.1038/nmeth.f.303. URL http://dx.doi.org/10.1038/nmeth.f.303.

[9] P. J. A. Cock, T. Antao, J. T. Chang, B. A. Chapman, C. J. Cox, A. Dalke, I. Friedberg, T. Hamelryck, F. Kauff, B. Wilczynski, and M. J. L. de Hoon. Biopython: freely available python tools for computational molecular biology and bioinformatics. Bioinformatics, 25(11):1422–1423, 2009. doi: 10.1093/bioinformatics/btp163. URL http://dx.doi.org/10.1093/bioinformatics/btp163.

[10] F. Sievers, A. Wilm, D. Dineen, T. J. Gibson, K. Karplus, W. Li, R. Lopez, H. McWilliam, M. Remmert, J. Soding, J. D. Thompson, and D. G. Higgins. Fast, scalable generation of high-quality protein multiple sequence alignments using clustal omega. Mol Syst Biol, 7(1):539–539, 2014. doi: 10.1038/msb.2011.75. URL http://dx.doi.org/10.1038/msb.2011.75.

